# A novel role of USP7 in imparting partial EMT state in colorectal cancer through the DDX3X-β-catenin axis

**DOI:** 10.1101/2023.01.11.523622

**Authors:** Bhaskar Basu, Subhajit Karmakar, Malini Basu, Mrinal K. Ghosh

## Abstract

Epithelial mesenchymal transition (EMT) is a fundamental and highly regulated process that is normally observed during embryonic development and tissue repair but is deregulated during advanced cancer. Classically, through the process of EMT, cancer cells gradually transition from a predominantly epithelial phenotype to a more invasive mesenchymal phenotype. Increasing studies have, however, brought into light the existence of unique intermediary states in EMT, often referred to as partial EMT states. Through our studies we have found the deubiquitinase USP7 to be strongly associated with the development of such a partial EMT state in colon cancer cells, characterized by the acquisition of mesenchymal characteristics but without the reduction in epithelial markers. We found USP7 to be overexpressed in colon adenocarcinomas and to be closely associated with advancing tumor stage. We found that functional inhibition or knockdown of USP7 is associated with a marked reduction in mesenchymal markers and in overall migration potential of cancer cells. Starting off with a proteomics-based approach we were able to identify and later on verify the DEAD box RNA helicase DDX3X to be an interacting partner of USP7. We then went on to show that USP7, through the stabilization of DDX3X, augments Wnt/β-catenin signaling, which has previously been shown to be greatly associated with colorectal cancer cell invasiveness. Our results strongly suggest a positive role of USP7 in the development of a partial mesenchymal phenotype in colorectal cancer.

## Introduction

From a global perspective colorectal cancer (CRC) is the third most diagnosed cancer and the second most in terms of mortality. An estimated 1.93 million new CRC cases were diagnosed, and 0.94 million CRC-related deaths were recorded globally in the year 2020. The number of new CRC cases recorded globally per year is also expected to increase to 3.2 million by the year 2040 [1]. About 25% of all CRC diagnoses generally develop into metastatic disease [2]. In such cases, conventional modes of treatment such as surgery and cytotoxic chemotherapies show little to no effect in halting the progression of disease and deterioration of patient health. For such reasons, a better understanding of the key molecular and cellular regulatory systems that drive progression of primary tumors towards metastasis is required.

Epithelial mesenchymal transition (EMT) is a naturally occurring cellular process that involves the transition of epithelial cells to their mesenchymal counterparts. A large number of investigations have demonstrated that EMT is a key phase for advanced stage cancers and is necessary for the onset of metastasis. During this phase cancer cells shed their epithelial characteristics and acquire a more mesenchymal and invasive phenotype. This transition is characterized by the repression of epithelial markers such as E-cadherin, occludin and claudin proteins and increased expression of mesenchymal markers such as N-cadherin, Vimentin and fibronectin [3, 4]. The regulation of EMT is facilitated by several core EMT transcription factors such as Snail, Slug, Twist, ZEB1 and ZEB2 and by many oncogenic signaling pathways such TGF-β, Wnt/β-catenin, Notch, Hedgehog and others [3, 5]. Recently it has been found that the process of EMT is not a binary one, and instead includes several unique intermediate states that are hybrids of epithelial and mesenchymal phenotypes and are termed partial EMT. Specific tumor subpopulations exhibiting such partial EMT have been identified and have also been shown to exhibit greater motility, chemoresistance and tumor initiating abilities compared to those tumor subpopulations that have undergone complete EMT [6–8].

DDX3 belongs to the DEAD-box RNA helicase family and consists of 2 homologs – DDX3X and DDX3Y [9]. DDX3X is ubiquitously expressed in most tissues. DDX3X has been shown to be involved in multiple modes of regulation of gene expression including transcription, pre-mRNA splicing, mRNA export and translation, and aberrant expression or mutations in DDX3X have been reported in many cancers [9–11]. DDX3X has been shown to be overexpressed in different cancers such as hepatocellular carcinoma, aggressive breast cancers and even colorectal cancer [10–12]. DDX3X has previously been shown to be involved in oncogenic transformation as its depletion is associated with G1/S arrest [13]. Most notably DDX3X has been shown to be implicated in the induction of EMT through the upregulation of Wnt/β-catenin signaling. DDX3X has been shown to upregulate β-catenin levels *via* mediators such as CK1ε and Rac1, which in turn leads to the activation of EMT and progression towards metastasis [14, 15].

The ubiquitin specific peptidase 7 (USP7) is one of the most extensively studied deubiquitinases and has been shown to act upon a rather vast substrate spectrum, thereby affecting various homeostatic and disease processes [16]. USP7 is most notably famous for its involvement in a dynamic relationship with the 2 proteins, p53 and MDM2, wherein it was revealed that USP7 preferentially acts on MDM2 during onset of cancers [17, 18]. Later it was established that both proteins interact with USP7, however MDM2 exhibits nearly ten-fold greater affinity for USP7 as compared to p53 [19, 20]. A similar relationship was shown to exist between Rb, MDM2 and USP7 [21]. Apart from these USP7 has been shown to interact with and stabilize many oncogenic substrates such as C-Myc, N-Myc, Suv39H1, Trip12, Ci/Gli, Notch 1and others [16, 22]. USP7 has been shown to be overexpressed in different cancers and has been shown to assume different regulatory roles during cancer progression in different tumors. The role of USP7 in the regulation of EMT and metastasis has yet to be properly assessed. It is possible that the stabilization of MDM2 by USP7 and subsequent ubiquitination-dependent degradation of p53 by MDM2 may lead to EMT induction as p53 plays an inhibitory role in EMT. However, the greatest obstacle to this theory is that MDM2 is also involved in the degradation of Slug, a key EMT transcription factor [23, 24].

In this study, we found that USP7 is overexpressed in CRCs and the increase in expression is associated with increasing tumor stage. CRC tissues displaying medium to high expression of USP7 also exhibited greater expression of the mesenchymal marker proteins N-cadherin and Vimentin. However, expression of epithelial marker protein E-cadherin showed little correlation with USP7. Inhibition of USP7 activity led to a marked reduction in migration ability of colon cancer cells and loss of mesenchymal identity. By utilizing a proteomic-based approach, we were able to identify DDX3X to be an interacting partner of USP7. At the mechanistic level, we evaluated that USP7 binds DDX3X and stabilizes it against degradation by the ubiquitin-proteasome system (UPS).

DDX3X in turn augments Wnt/β-catenin signaling to promote the development of partial EMT state. Our results demonstrate the role of USP7 in the development of a partial EMT state through DDX3X-mediated augmentation of Wnt/β-catenin signaling.

## Materials and methods

### Bioinformatics analysis

USP7 mRNA expression data from the TCGA database was analyzed and plotted against 24 different cancer types and data normalization in RNA expression profile was done using UALCAN (http://ualcan.path.uab.edu/analysis.html [25]. We applied a cut-off value of p-0.05 as significant with a fold change boundary within 2. The expression of USP7 in different datasets of colorectal carcinoma was further checked using the online Oncomine platform [26]. The specimens were compared as cancer vs. normal patients. The Gene Expression Profiling Interactive Analysis (GEPIA) database (http://gepia.cancer-pku.cn/) was used to evaluate to the differential expression of USP7 in COAD tissues vs normal tissues [27].

### Human colorectal cancer tissues

Formalin-fixed paraffin-embedded sections of stage 3 CRC tissues (n=20), derived from post-surgical humans collected from patients of West Bengal, India was used in this study. The samples were collected in accordance with all medical and ethical regulations, including patient consent, and with formal approval from the institutional ethical committees of both CSIR-IICB and Park Clinic.

### Immunohistochemistry (IHC) and analysis

Immunohistochemical studies were conducted as described before [28]. Staining scores were defined as the cell staining intensity (0 = nil; 1 = weak; 2 = moderate; and 3 = strong) multiplied by the respective percentage of labeled cells (0–100%), leading to H-scores falling within the range 0 to 300. An H-score above 150 was rated as “high expression”, between 100-150 was rated as “medium expression”, and below 100 was rated as “low expression”. Images were captured at 60X magnification by EVOS XL Cell Imaging System (Life Technologies-Thermo Fischer Scientific).

### Cell culture, transfection and treatments

HCT116 cells were maintained in McCoy’s 5A media while HEK293, and SW480 cells were cultured in Dulbecco’s modified eagle medium (DMEM) supplemented with 10% heat inactivated fetal bovine serum, and antibiotics penicillin, streptomycin and gentamycin at previously described doses [29]. Cells were cultured at 37°C in a humid incubator with a set atmosphere of 5% CO_2_. Transfection of cells was carried out using Lipofectamine 3000 (Invitrogen) in accordance with the manufacturer’s instructions. siRNA mediated silencing of USP7 was carried out using Lipofectamine RNAimax (Invitrogen). Cells were harvested 48 h following transfection.

DNA, transfections were done at least 24 h prior to drug treatment. The USP7 inhibitors P5091 and P22077 (Sigma Aldrich) were administered at 10μM and 15μM respectively for 24 and 48 h time periods. Cyclohexamide (Sigma Aldrich) was administered at 50μg/ml for indicated time periods. MG132 (Sigma Aldrich) was administered at 40μM for 4 h.

### Plasmids and small interfering RNA (siRNA)

Plasmids expressing FLAG-tagged USP7 (empty vector: pIRES-hrGFP-1a), GFP-tagged USP7 (empty vector: pGZ-21dx), and USP7 catalytic mutant (C223S), and GFP-tagged USP7 deletion mutants (ΔTRAF, ΔUBL, and RING only) were described previously [21]. FLAG-tagged DDX3X (empty vector: pCS2-N-FLAG) was a gift from Dr. Beat Suter (University of Bern, Switzerland) [30]. HA-tagged ubiquitin (empty vector: pRK5-HA) was obtained from ADDGENE (17608). Control siRNA and siRNA targeting USP7 was obtained from Santa Cruz Biotechnology and added at a final concentration of 30 picomolar.

### Western blotting

Cell lysates were prepared by resuspension of cells in whole cell lysis buffer (10 mM Tris, 150 mM NaCl, 1% deoxycholate, 1 mM EDTA, 0.1% SDS, 1% NP-40, 1 mM sodium orthovanadate) supplemented with protease inhibitor cocktail (Calbiochem). Protein concentration was measured by Bradford method. Total 40 μg of protein was used for analysis. Proteins were separated *via* SDS-PAGE and transferred to a PVDF membrane. Membranes were blocked with 5% BSA in TBS-T and incubated with indicated primary antibodies diluted in 1% BSA in TBS-T overnight at 4°C. On the following day, membranes were incubated with respective HRP-conjugated secondary antibodies (Sigma Aldrich) for 2 h at room temperature, membrane was developed using Classico ECL Millipore chemiluminescence substrate. Visualized images were captured using Bio-Rad Chemi-doc. Primary antibodies against the following target proteins were used in this investigation: N-cadherin, E-cadherin, DDX3X (Abclonal), Vimentin (Cell Signaling Technology), USP7, β-catenin, GFP (Santa Cruz Biotechnology), Flag, HA, Actin (Sigma Aldrich). Densitometric quantification of specific blots was performed using ImageJ. Densitometric values for immunoblots were calculated with respect to the respective loading control.

### GST-pulldown coupled to mass spectrometry

HEK293 cells were grown in 100 mm cell culture plates up to 90% confluency, harvested and lysates were prepared in a similar manner as that described in the immunoprecipitation section. Lysates were equally separated into two samples. Purified GST-USP7 protein was added to one of the samples while purified GST protein was added to the other. Mixtures were incubated overnight at 4 °C with gentle rotation. Glutathione coated beads were added to the mixtures on the next day and incubated for 3 h at 4 °C, following which beads were separated out through gentle centrifugation. Beads were washed 3 times with excess GST lysis buffer (50 mm Tris/HCl, pH 7.5; 300 mm NaCl; 1 mm EDTA;1% Triton-X-100; pH 8) supplemented with protease inhibitor cocktail. Bead bound proteins were eluted by boiling in 2X protein gel loading buffer and separated using SDS-PAGE. Proteins in the gel were silver stained, each protein lane of the gel was divided in to 3 sections (upper, middle and lower), gel sections were processed, the proteins were trypsinized in gel, and the peptides were extracted and prepared for mass spectrometry following a previously described protocol [31]. Peptide signatures were analyzed by liquid chromatography-tandem mass spectrometry (LC-MS/MS) using a LTQ Orbitrap system (Thermo) and identified using Mascot software (Matrix Science, United Kingdom).

### Immunoprecipitation

Cells were resuspended in RIPA buffer supplemented with 2X protease inhibitor cocktail (Calbiochem), sonicated and centrifuged at 12000g for 15 minutes to obtain lysates. Lysates were pre-cleared by incubating with protein A or G sepharose beads (GE Healthcare) for 1 h at 4 °C. Equal amounts of lysates were then incubated with antibodies of interest overnight at 4 °C. The immunocomplexes were harvested by incubating with protein A or G sepharose beads, with gentle rotation for 4 h at 4 °C. Beads were collected by centrifugation, were washed with excess RIPA buffer and boiled in 2X protein gel loading buffer to elute proteins before running SDS PAGE followed by Western blotting. For reference, 3% input was run separately and probed with indicated antibodies.

### Protein-protein docking

The 3D crystal structures of USP7 TRAF-like domain of HAUSP/USP7 (PDB id: 2F1W), and human DEAD-box RNA helicase DDX3X protein (PDB id: 2I4I) were extracted from protein databank (PDB). To obtain a docked structure and a stable docked pose, TRAF domain of USP7 was docked with DDX3X using clusPro, PatchDock and SwarmDock docking programs, respectively [33–35]. Top 20 docking solutions for each pair were selected based on docking score and used for RMSD based clustering using UCSF Chimera [36]. The ranking of the largest cluster was done based on the average docking score of the docking solutions present in the largest cluster.

Top 3 pairs based on highest rank were chosen from each of the docking programs and compared. These pairs were selected for visualizing as the probable interacting domains using PyMol (https://www.schrodinger.com/products/pymol). The stereo-chemical properties of the highest docked model and the fold compatibility of the final models (USP7 and DDX3X) were validated by Rampage28, Prove29, Verify3D30, and Errat31 programs. The USP7-DDX3X complexes after docking were validated by estimating the true-like complex probability at PCPIP (Protein Complex Prediction by Interface Properties) server. Electrostatics calculations were done using APBS (Adaptive Poisson-Boltzmann Solver) tool [37] after assigning the atomic charges by PDB2PQR (https://server.poissonboltzmann.org), and the electrostatic potential map obtained was visualized using 3Dmol (http://3dmol.csb.pitt.edu/).

### Deubiquitination assay

Deubiquitination assay was performed under denaturing conditions. Cells were briefly treated with the indicated amount of MG-132 for 4 h before harvesting to allow for the accumulation of poly-Ub proteins. Cell lysates were prepared using RIPA buffer supplemented with 10 mM NEM (Sigma) and protease inhibitor cocktail. The lysates were immunoprecipitated with indicated antibodies and probed with anti-HA antibodies (Sigma Aldrich).

### Immunofluorescence microscopy

HCT116 cells were seeded on coverslips placed in 35 mm culture dishes and grown until ~ 40% confluency. Cells were then fixed in 4% paraformaldehyde, permeabilized with 0.5% Triton X-100, blocked with 5% BSA in PBS, washed and then immunostained with primary antibodies for USP7 (Santa Cruz Biotechnology) and DDX3X (Abclonal) and secondary antibodies conjugated with the following fluorophores – Alexafluor 488 (green) and Alexafluor 568 (red). Cell nuclei were counterstained with Hoechst. Images were captured at 60X and 120X optical zoom using the FluoView FV10i confocal laser scanning microscope (Olympus Life Science). Quantification of fluorescence colocalization was carried out using ImageJ.

### In vitro scratch/ wound healing assay

Cells were seeded at the recommended density into 60 mm culture dishes so that they were nearly at 80-90% confluency after 24 h. Following this, fine but detectable scratches were made through the cells using sterile tips. Cells were observed under 10X objective at indicated time points to assess the extent of migration through the scratches.

### Transwell migration assay

A 6-well hanging transwell cell culture insert (Millipore) with 8.0 μm pore size PET membrane was used for the migration assay. HCT116 cells at a density of 5 x 10^5^ cells in 500 μl of serum free media were seeded in the upper chamber. The lower chamber was filled with 1 ml of complete media as a chemoattractant. Following incubation for 24 h at 37°C, cells remaining in the upper chamber were removed using a cotton swab. Cells attached to the lower surface of the membrane represented migrated cells and were first fixed in 3.7% formaldehyde in PBS, followed by permeabilization in methanol and stained using crystal violet solution. Photographs of migrated cells were taken at 20x optical zoom from 3 different fields to assess changes in migratory behavior.

### Statistical analysis

Statistical significance in case of quantification of wound healing assay was drawn by conducting two-way Annova. For the remainder of experiments statistical significance was drawn by conducting Two Samples Student’s t-test. The significance values were represented as * P < 0.05, **P ≤ 0.01, and ****P ≤ 0.0001. All statistical analyses were performed using either SPSS (IBM) or the GRAPHPAD PRISM 8.4 software.

## Results

### USP7 is overexpressed in Pan-cancer and colorectal cancer

To explore the role of USP7 in cancer we first analyzed USP7 mRNA expression data of 24 different cancers available in the TCGA database using the UALCAN online web tool (Fig. 1A). USP7 expression was upregulated in 17 out of the 24 cancer types. Of these, the results of colon adenocarcinoma (COAD) were of particular interest to us, both for its global relevance and its rising prevalence amidst the Indian cancer scenario. Using the COAD dataset, we further assessed USP7 mRNA expression in normal sample vs primary tumor (Fig. 1B). USP*7* overexpression was also detected in primary colon cancer samples compared to normal samples. Oncomine analysis of Hong colorectal (Fig. 1C) and Skrzypczak colorectal datasets (Fig. S1A) also revealed overexpression of USP7 in CRC samples compared to normal colon tissue. USP7 overexpression was associated with increasing tumor stage, with stage 3 tumors showing the highest USP7 expression (Fig. S1B). Differential expression of USP7 in COAD tissues and normal tissues was further evaluated using GEPIA (Fig. 1D). Finally, USP7 protein expression data from the CPTAC database consisting of 97 CRC tissue samples and 100 adjacent normal tissue samples further validated the mRNA expression results (Fig. 1E).

**Fig 1.**
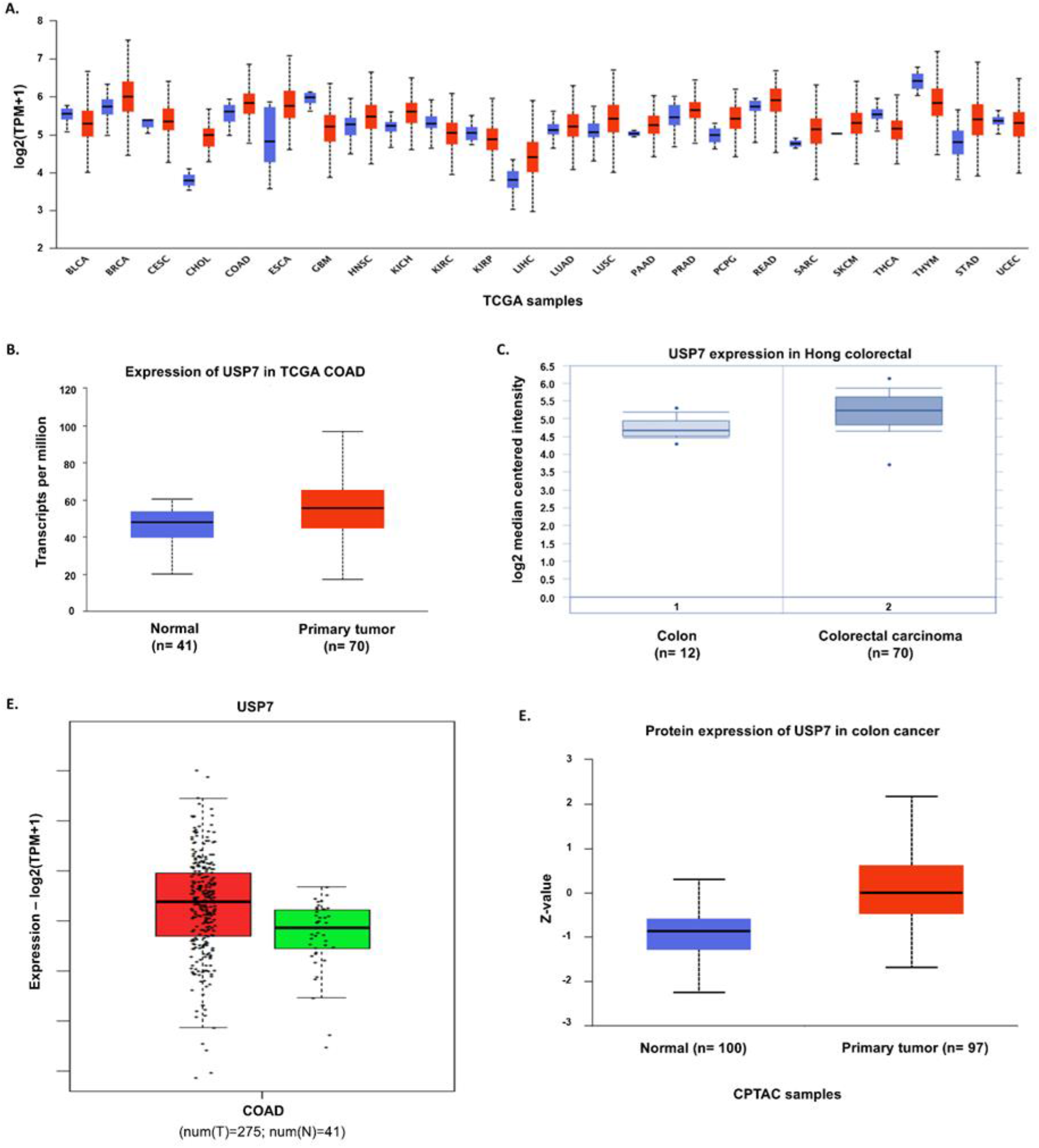
USP7 is overexpressed in colon cancers. (A) Pan-cancer analysis of USP7 expression across different cancers using the UALCAN platform. BLCA, bladder urothelial carcinoma; BRCA, breast invasive carcinoma; CESC, cervical squamous cell carcinoma and endocervical adenocarcinoma; CHOL, cholangiocarcinoma; COAD, colon adenocarcinoma; ESCA, esophageal carcinoma; GBM, glioblastoma multiforme; HNSC, head and neck squamous cell carcinoma; KICH, Kidney chromophobe; KIRC, kidney renal clear cell carcinoma; KIRP, kidney renal papillary cell carcinoma; LIHC, liver hepatocellular carcinoma; LUAD, lung adenocarcinoma; LUSC, lung squamous cell carcinoma; PAAD, pancreatic adenocarcinoma; PRAD, prostate adenocarcinoma; PCPG, pheochromocytoma and paraganglioma; READ, rectum adenocarcinoma; SARC, sarcoma; SKCM, skin cutaneous melanoma; THCA, thyroid carcinoma; THYM, thymoma; STAD, stomach adenocarcinoma; UCEC, uterine corpus endometrial carcinoma. Blue boxes represent normal tissue groups while red boxes represent cancer tissue groups. (B) USP7 gene expression level in in normal colon vs primary tumor tissues from the TCGA dataset. (C) USP7 gene expression in normal colon vs colon carcinoma tissues from the Hong colorectal dataset. (D) Gene expression profiling interactive analysis (GEPIA) of USP7 in COAD samples (P < 0.05). Red box represents tumor tissue group (T) and green box represents adjacent normal tissue group (N). (E) USP7 protein expression in the primary tumors and adjacent normal tissues of CRC patients from CPTAC database.

### USP7 expression is associated with increased mesenchymal features in CRC

EMT is a characteristic feature of advanced stage tumors and is essential for migration of cancer cells from the primary tumor site into adjacent tissues and lymph nodes and is a preceding step before metastasis. Because of the clear upregulation of USP7 in colon cancers, we decided to explore possible associations between USP7 expression and EMT in CRC. We performed immunohistochemical analysis to assess the expression of USP7, the 2 mesenchymal markers N-cadherin and Vimentin, and the epithelial marker E-cadherin in 20 CRC tissue samples (Fig. 2A).

**Fig 2.**
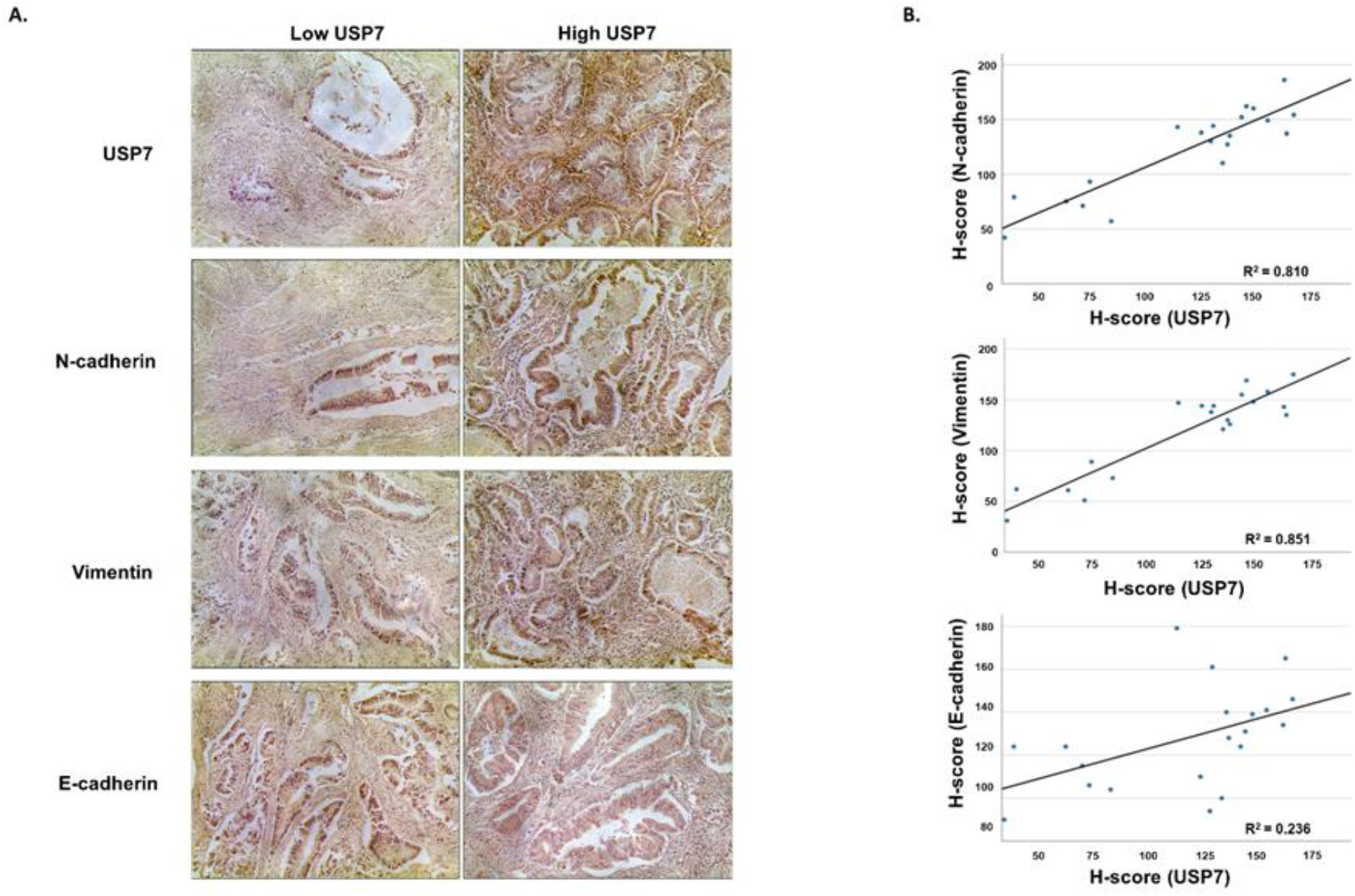
USP7 expression is correlated with increased mesenchymal features in CRC. (A) Representative immunostaining results of USP7, N-cadherin, Vimentin and E-cadherin in colorectal tumors. (B) Depiction of correlation (R^2^ linear) between USP7 and indicated EMT markers estimated from IHC images of colorectal tumors.

High N-cadherin and Vimentin expression was observed in tumors exhibiting medium to high USP7 expression compared to tumors exhibiting low USP7 expression. Interestingly, E-cadherin expression showed very little correlation with USP7 expression and mostly displayed medium staining intensity throughout the entire set of CRC tissue samples (Fig. 2B). This observation was in line with previous studies where it has been reported that high percentage of CRC primary tumors and even lymph nodes showed E-cadherin staining [32, 33]. A summarization of the immunohistochemical analysis has been provided in Table 1. Taken together, we found strong correlation between USP7 and mesenchymal marker expression in CRC. So, we decided to investigate further into the possible role USP7 may play in advancing CRC progression to metastasis.

**Table 1.** Summary of immunohistochemical expression of USP7 and select EMT markers in CRC tissues (n = 20).

|  | N-cadherin |  |  |  | Vimentin |  |  |  | E-cadherin |  |  |  |
| --- | --- | --- | --- | --- | --- | --- | --- | --- | --- | --- | --- | --- |
| USP7 | Low | Medium | High | Total | Low | Medium | High | Total | Low | Medium | High | Total |
| Low | 6 | 0 | 0 | 6 | 6 | 0 | 0 | 6 | 1 | 5 | 0 | 6 |
| Medium | 0 | 7 | 3 | 10 | 0 | 8 | 2 | 10 | 1 | 7 | 2 | 10 |
| High | 0 | 2 | 2 | 4 | 0 | 2 | 2 | 4 | 0 | 3 | 1 | 4 |
| Total | 6 | 9 | 5 | 20 | 6 | 10 | 4 | 20 | 2 | 15 | 3 | 20 |

H-score below 100 is considered as Low expression, between 100-150 as medium expression, and above 150 is considered as High expression.

### USP7 inhibition/knockdown is associated with repression of mesenchymal phenotype and reduced migration in colon cancer cells

In order to build a better understanding of the possible function of USP7 in controlling colon cancer migration, we performed a wound healing assay utilizing the HCT116 colon cancer cell line. We observed that the migratory ability of HCT116 cells was significantly impaired upon prolonged exposure to either of the potent USP7 inhibitors P5091 and P22077 as compared to DMSO control (Fig. 3A). Additionally, a transwell migration assay carried out using the same cell line revealed that USP7 inhibition significantly reduced the migration ability of the cell line (Fig. 3B). These results suggested that USP7 positively regulates cellular migration and that its functional inhibition was detrimental towards the overall migration potential of colon cancer cells. In order to elaborate on these findings, we decided to assess the status of mesenchymal marker proteins upon USP7 inhibition. Both the colon cancer cell lines, HCT116 and SW480, showed reduction in the level of the mesenchymal markers N-cadherin and Vimentin upon exposure to either P5091 or P22077 (Fig. 3C). This effect appeared to be much more apparent in case of HCT116 cells compared to SW480. We believe that this was most likely due to HCT116 cells displaying greater levels of endogenous USP7 than SW480 cells (Fig. S2A), thereby exhibiting a greater reliance on the protein. A subtle rise in the levels of the epithelial marker E-cadherin was also observed in both the cell lines upon exposure to USP7 inhibitors. Similar observations were documented upon exposure of the HT29 cell line to P5091 (Fig.S2B)

**Fig 3.**
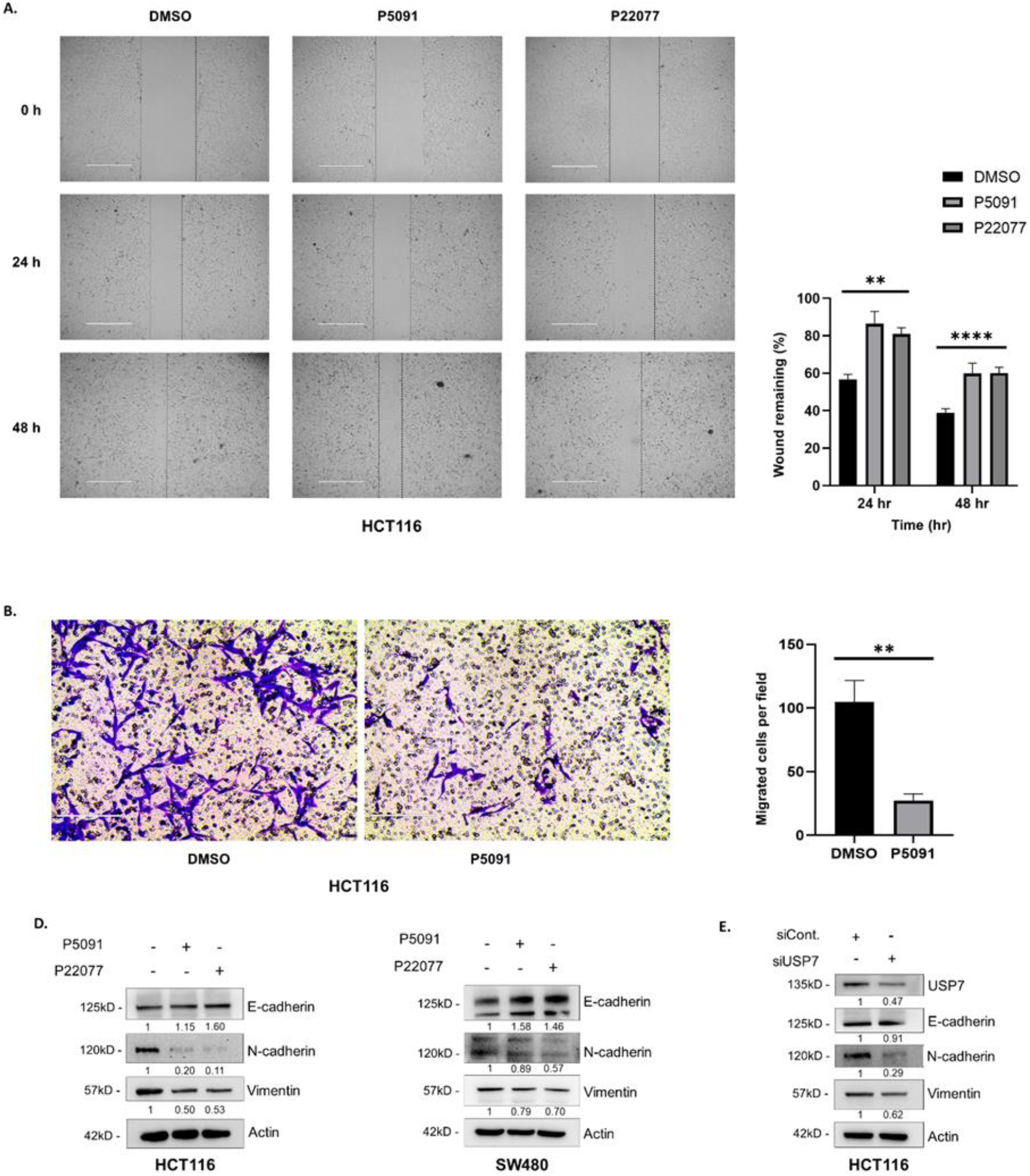
USP7 inhibition/knockdown is associated with repression of mesenchymal phenotype and reduced migration in colon cancer cells. (A) HCT116 cells were treated with either DMSO control, P5091 (10 μM) or P22077 (15 μM) and then subjected wound healing assay. Pictures were taken at 0, 24, and 48 h. (B) Bar graphs represent percentage of gap/wound remaining at indicated time points following generation of the scratch. (C) HCT116 cells were treated with either DMSO control or P5091 (10 μM) and then subjected to transwell migration assay. Pictures were taken post 24 h of incubation. (D) Bar graphs represent degree of cellular migration following 24 h of treatment. (E) HCT116 (right panel) and SW480 cells (left panel) were treated with either DMSO control, P5091 (10 μM) or P22077 (15 μM) for 48 h. Whole cell lysates were analyzed by western blotting using the indicated primary antibodies. (**F**) USP7 silencing was carried out using small interfering RNA (siRNA). Whole cell lysates were analyzed by western blotting using specific primary antibodies. Error bars in all the indicated subfigures represent mean (+) s.d. from three independent biological repeats. Indicated P values were calculated using two-way Annova (2B) or Student’s t-test (2D). P ≤ 0.01 is represented as ** and P ≤ 0.0001 as ****.

In order to rule out any off-target effects of the 2 inhibitors and to decisively conclude if the repression of mesenchymal phenotype was solely due to USP7 inhibition, we depleted endogenous USP7 in HCT116 cells by using USP7-targeting small interfering RNA (siRNA). A similar downregulation of the mesenchymal markers N-cadherin and Vimentin was observed which validated the results of our previous USP7 inhibition experiments (Fig.3D). E-cadherin levels, however, exhibited no significant changes due to USP7 knockdown. These findings are in line with the results of our immunohistochemistry analysis of colon cancer tissues. These data indicate that USP7 expression promotes the enhanced expression of mesenchymal markers but elicits minimal alteration in epithelial marker expression.

### Identification of novel USP7 interactors with links to EMT induction

To gain better understanding of the USP7 interactome and its possible connections to EMT, we incubated purified GST-USP7 protein with HEK293 lysates, following which pulldown was carried out using glutathione coated agarose beads. Recovered bound proteins were identified using mass spectrometry (Nano LC-MS/MS). To eliminate non-specific interactions total spectral counts of proteins recovered with GST-USP7 were compared with control sets where purified GST protein was used for incubation with lysates. The top USP7 interactions identified are listed in Table 2 and the complete interaction data is provided in Supplemental Table S1.

**Table 2.** GST pulldown-mass spectrometry results for GST-USP7.

| Gene name | Protein ID | Protein classification | GST-USP7 Score (spectral counts) | GST score (spectral counts) | Sequence coverage (%) | Molecular weight (kDa) |
| --- | --- | --- | --- | --- | --- | --- |
| USP7 | Q93009 | Deubiquitinase | 2024.79 | 0 | 69.15 | 128.2 |
| DDX3X | O00571 | RNA helicase | 203.35 | 0 | 21.3 | 73.1 |
| RUVBL1 | Q9Y265 | DNA helicase | 144.56 | 0 | 20.39 | 50.2 |
| RUVBL2 | Q9Y230 | DNA helicase | 126.1 | 0 | 17.71 | 51.1 |
| MCM5 | P33992 | DNA helicase complex subunit | 64.02 | 0 | 5.59 | 82.2 |
| DHX40 | Q8IX18 | RNA helicase | 35.27 | 0 | 1.67 | 88.5 |
| FLII | Q13045 | Transcriptional coactivator | 29.4 | 0 | 0.79 | 144.7 |

Consistent with previous reports, USP7 was shown to interact with the previously established DEAH box RNA helicase DHX40 [34]. In addition, several previously unreported interactions were also recorded such as the DNA helicases RUVBL1 and RUVBL2, the mini-chromosome complex (MCM) subunit MCM5, and the co-activator of hormone activated nuclear receptors FLII. However, what interested us the most was the presence of the DEAD-box RNA helicase DDX3X among the list of identified proteins. DDX3X has previously been shown to aid in EMT induction through the stabilization of β-catenin and augmentation of Wnt/β-catenin signalling through different intermediary proteins [14, 15]. As these functions of DDX3X were very much in line with our investigative goals, we decided to look further into the nature of the interaction between USP7 and DDX3X.

### USP7 colocalizes and interacts with DDX3X

In order to validate the results of our mass spectrometric analysis, we decided to inspect the interaction between USP7 and DDX3X *in vitro*. Immunofluorescence staining of both USP7 and DDX3X in HCT116 cells revealed that while USP7 was predominantly accumulated in the nucleus, DDX3X was localized in both the cytosol and the nucleus. As such maximum colocalization of the two proteins was observed in the nucleus (Fig. 4A&B). We transiently expressed GFP-tagged USP7 and Flag-tagged DDX3X in HEK293 cells followed by immunoprecipitation using antibody against GFP. Subsequent western blotting revealed FLAG tagged DDX3X to co-immunoprecipitate along with exogenous USP7 (Fig. 4C). We then performed a similar experiment wherein we transiently expressed Flag-tagged DDX3X in HEK293 cells followed by immunoprecipitation using antibodies against FLAG. Endogenous USP7 was observed to coimmunoprecipitate with exogenous DDX3X upon western blotting (Fig. 4D). These findings confirmed that USP7 does indeed interact with DDX3X and that the interaction occurs primarily in the nucleus.

**Fig. 4.**
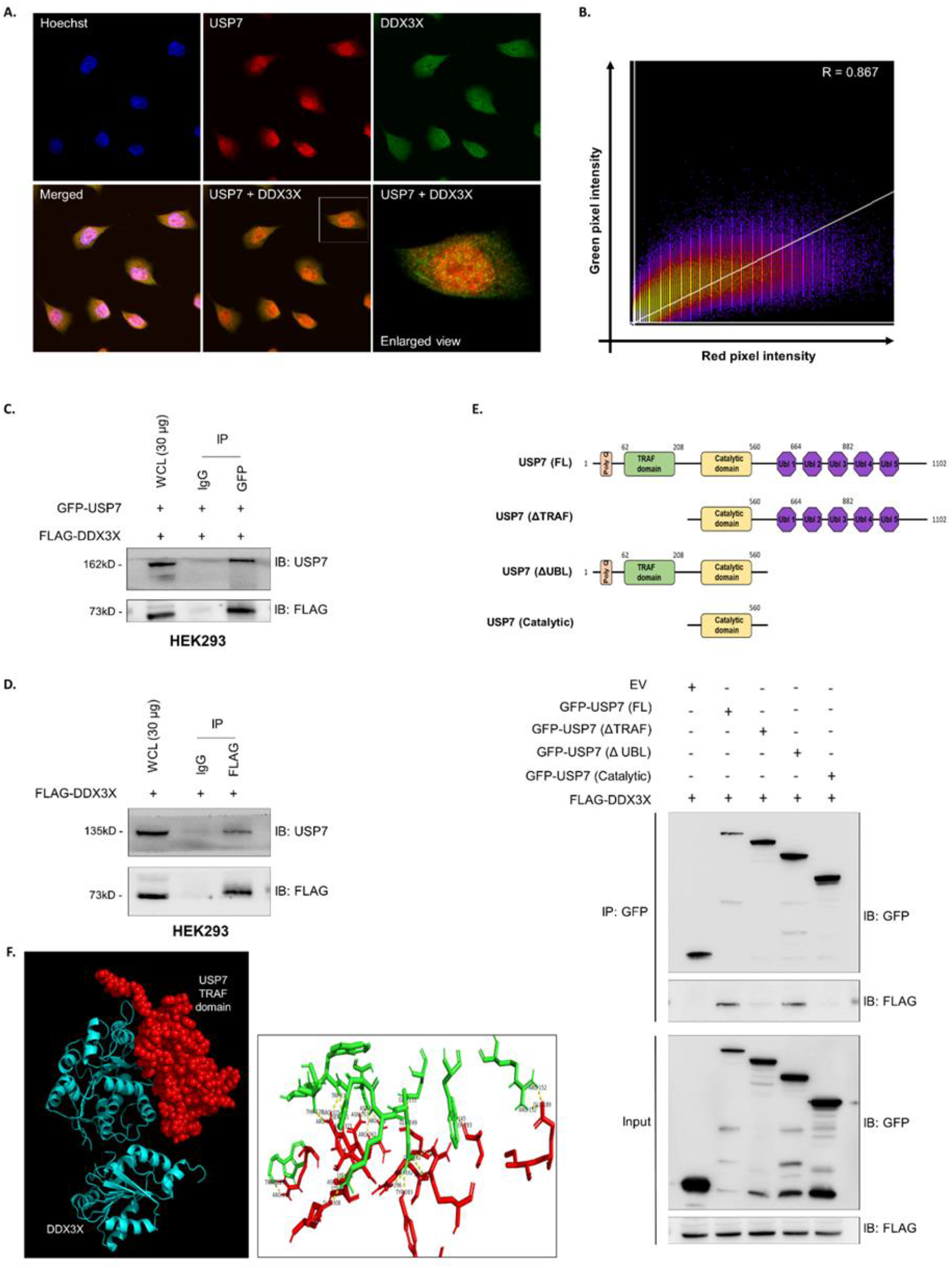
USP7 interacts with DDX3X. (A) HCT116 cells were fixed and immunostained using fluorophore tagged antibodies specific for USP7 (red) and DDX3X (green). Cell nuclei were counterstained with Hoechst (blue). Orange/yellow fluorescence indicates regions of colocalization of the 2 proteins. (B) Scatter plot of red vs green pixel intensities represents degree of colocalization of the 2 proteins. Pearson correlation coefficient, R = 0.867. (C) HEK293 cells were transfected with FLAG-DDX3X and full length GFP-USP. Equal amounts of whole cell lysate were used for immunoprecipitation using either GFP antibody or non-specific mouse IgG, followed by western blotting with indicated antibodies to assess coimmunoprecipitation status. (C) HEK293 cells were transfected with FLAG-DDX3X. Equal amounts of whole cell lysate were used for immunoprecipitation using either FLAG antibody or non-specific rabbit IgG, followed by western blotting with indicated antibodies to assess coimmunoprecipitation status. (D) Schematic representation of USP7 deletion mutants used for domain mapping. (E) HEK293 cells were transiently transfected with FLAG-DDX3X and full length (FL) GFP-USP or with the GFP-tagged USP7 deletion mutants. Immunoprecipitation carried out with equal amounts of whole cell lysate and by using antibody against GFP. Western blotting was carried out using the indicated antibodies. (F) Protein-protein docking model of the USP7 TRAF domain in complex with DDX3X (Left panel). USP7 TRAF domain is represented as space filling model (red) while DDX3X is represented as ribbon diagram (blue). Stereo view of predicted interactions between USP7 TRAF domain and DDX3X (Right panel). Dashed yellow lines represent hydrogen bonds.

USP7 consists of 4 structural domains – an N-terminal poly-glutamine stretch (poly Q), the TRAF (TNF receptor associated factor) domain, the catalytic (CAT) domain and a C-terminal domain (CTD) [16, 19]. Majority of USP7-substrate interactions are mediated by means of the N-terminal TRAF domain [19, 20, 35]. In order to gain better insight into the mode of interaction between USP7 and DDX3X, we re-performed the USP7 co-IP experiments in HEK293 cells using either full length (FL) GFP-tagged USP7 or a set of GFP-tagged USP7 deletion mutants that lacked either the TRAF domain (ΔTRAF) or the UBL domains (ΔUBL), or consisted of only the catalytic domain (Fig. 4E). Of these only full length USP7 and the ΔUBL mutants were able to interact with DDX3X, indicating that the interaction between USP7 and DDX3X required the presence of the USP7TRAF domain (Fig. 4E). In addition, examination of the amino acid sequence of DDX3X revealed the presence of four putative USP7 TRAF domain binding motifs (P/AXXS) [19] (Fig. S3A).

Protein-protein docking using ClusPro, a web-based server which uses a PIPER based Fast Fourier Transform (FFT) correlation approach along with root-mean-square deviation (RMSD) based clustering of the 1000 lowest energy structures generated, demonstrated a rigid body docking. We then found out the largest clusters (cluster 0) was the best suited docked models with significant lowest energy weighted score of −982.2 and center weighted score of −921.9. Docked model.000.00 of USP7-TRAF domain and DDX3X domain protein interaction was found to display the strongest protein-protein interaction which was then visualized by PyMol (Fig. 4F). We also demonstrated the possible amino acids interaction of the best suited docked structure (Fig. 4F). Next, we assessed the quality of the docked structure of USP7-TRAF and DDX3X, by accessing the structure with SAVES v6.0, assessment tool and found that the docked structure gave an ERRAT score of about 92.50 and also 81.9% residues were detected in the most favored region of the Ramachandran plot (Fig. S3B).

The Adaptive Poisson–Boltzmann Solver (APBS) software uses force field parameters such as atomic charges and radii. Default APBS charges and atomic radii were used, and the electrostatic potential was measured in eV, with a range of surface potential from 5kT/e to −5kT/e that depicts the overall property of protein model along the three spatial dimensions where negatively and positively charged surface areas are depicted in red and blue, respectively (Fig. S3C & D).

### USP7 protects DDX3X from ubiquitination-dependent proteasomal degradation

USP7 being a deubiquitinase has been shown to deubiquinate and protect many of its interacting partners from ubiquitination-dependent proteasomal degradation. Hence, it was highly likely that USP7 may serve a protective role for DDX3X. We found that in HCT116 cells upon USP7 inhibition a marked reduction in DDX3X levels was incurred (Fig. 5A). Transient expression of exogenous USP7 in HEK293 was able to significantly increase endogenous DDX3X levels (Fig. 5B). Similarly, over-expression of USP7 was able to elicit an observable increase in levels of exogenous DDX3X and this increase was proportional to the amount of USP7 DNA transfected (Fig. 5B & 5C). Interestingly, the catalytically inactive USP7 mutant (C223S) was unable to elicit a similar positive response (Fig. 5D). This suggested that the upregulation in DDX3X protein levels upon USP7 overexpression was dependent on the deubiquitinase activity of USP7.

**Fig. 5.**
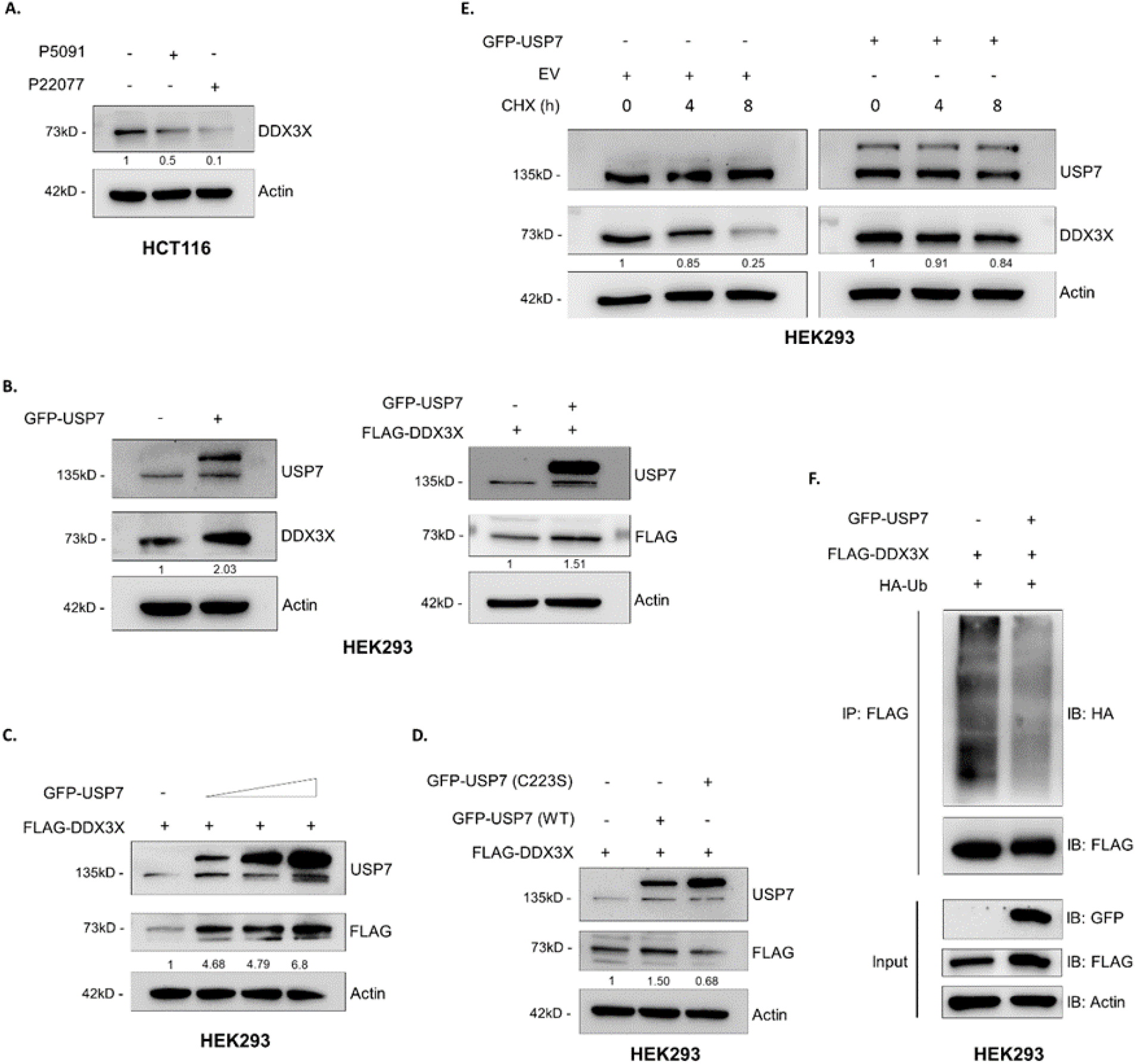
USP7 protects DDX3X from ubiquitination-dependent proteasomal degradation. (A) HCT116 cells were treated with either DMSO control, P5091 (10 μM) or P22077 (15 μM) for 48 h. Whole cell lysates were analyzed by western blotting using the indicated primary antibodies. (B) On the right panel, HEK293 cells were transfected with either empty vector or GFP-USP7. Whole cell lysates were analyzed by western blotting using indicated primary antibodies. On the left panel, HEK293 cells were transfected with FLAG-DDX3X and either empty vector or GFP-USP7. Whole cell lysates were analyzed by western blotting using indicated primary antibodies. (C) HEK293 cells were transfected with FLAG-DDX3X and either empty vector GFP-USP7 (1 μg, 1.5 μg, and 2 μg DNA transfected). Whole cell lysates were analyzed by western blotting using indicated primary antibodies. (D) HEK293 cells were transfected with FLAG-DDX3X and either empty vector, GFP-USP7 or GFP-USP7 (C223S). Whole cell lysates were analyzed by western blotting using indicated primary antibodies. (E) HEK293 cells were transfected with either empty vector or GFP-USP7. Cyclohexamide (CHX, 50 μg/ml) treatment was carried out for indicated time periods. Whole cell lysates were analyzed by western blotting using indicated primary antibodies. (F) *In vitro* deubiquitination assay. HEK293 cells were transfected with FLAG-DDX3X, HA-Ub and either empty vector or GFP-USP7. This was followed by MG132 (30 μM) treatment for 6 h. Immunoprecipitation was carried out with equal amounts of whole cell lysate, using antibodies against FLAG. Western blotting was carried out using the indicated antibodies.

Regulation of DDX3X protein stability by the UPS has not received much attention yet. So far, only the E3 ligase RNF39 has been shown to mediate the K48-linked polyubiquitination and proteasomal degradation of DDX3X [36]. Exogenously expressed USP7 was able to significantly extend the half-life of DDX3X as compared to empty vector control (Fig. 5E). In addition, transient expression of USP7 in HEK293 cells was able to elicit a marked reduction in the levels of ubiquitinated forms of DDX3X (Fig. 5F). This verified the observed upregulation of DDX3X in response to USP7 overexpression involved the deubiquitination of ubiquitinated DDX3X, thereby leading to its increased stability. Our findings strongly indicate that USP7 is a novel deubiquitinase essential for the stabilization of the RNA helicase DDX3X.

### USP7 augments Wnt/β-catenin signaling through DDX3X stabilization to promote mesenchymal phenotype

DDX3X has consistently been reported to positively regulate Wnt/β-catenin signaling, and thereby promote EMT and cancer cell migration [14, 15]. Up till now the relationship of USP7 with Wnt/β-catenin signaling has yet to reach a decisive conclusion. Whereas the majority of studies have reported a positive regulatory role of USP7 in Wnt/β-catenin signaling [37–39] another study has shown that USP7 actually inhibits Wnt/β-catenin signaling by stabilizing Axin, a key scaffolding protein involved in the formation of the β-catenin destruction complex [40]. Keeping all of this in mind we decided to investigate whether USP7 can positively regulate Wnt/β-catenin signaling through DDX3X stabilization.

Firstly, we checked the effect of USP7 inhibition on β-catenin levels. Treatment of HCT116 cells with either P5091 or P22077 produced a significant reduction in the levels of β-catenin protein and was proportional to the reduction in levels of DDX3X protein (Fig. 6A). The reduction in β-catenin levels upon USP7 inhibitor treatment was nullified upon short exposure with the proteasome inhibitor MG132. This suggested that the decrease in β-catenin protein levels upon USP7 inhibition was due to disruption at the protein level rather than at the gene expression level (Fig 6B). We also found that USP7 inhibition elicited a greater decrease in nuclear β-catenin levels compared to its cytosolic version (Fig. 6C). Exogenous expression of DDX3X in HCT116 cells was able to produce a marked increase in the level of β-catenin and was also able to partially rescue β-catenin levels even after exposure of HCT116 cells to P5091 (Fig. 6D). Again, this was in line with previously reported findings that DDX3X is a major controller of β-catenin and maintains β-catenin at the posttranslational level.

**Fig. 6.**
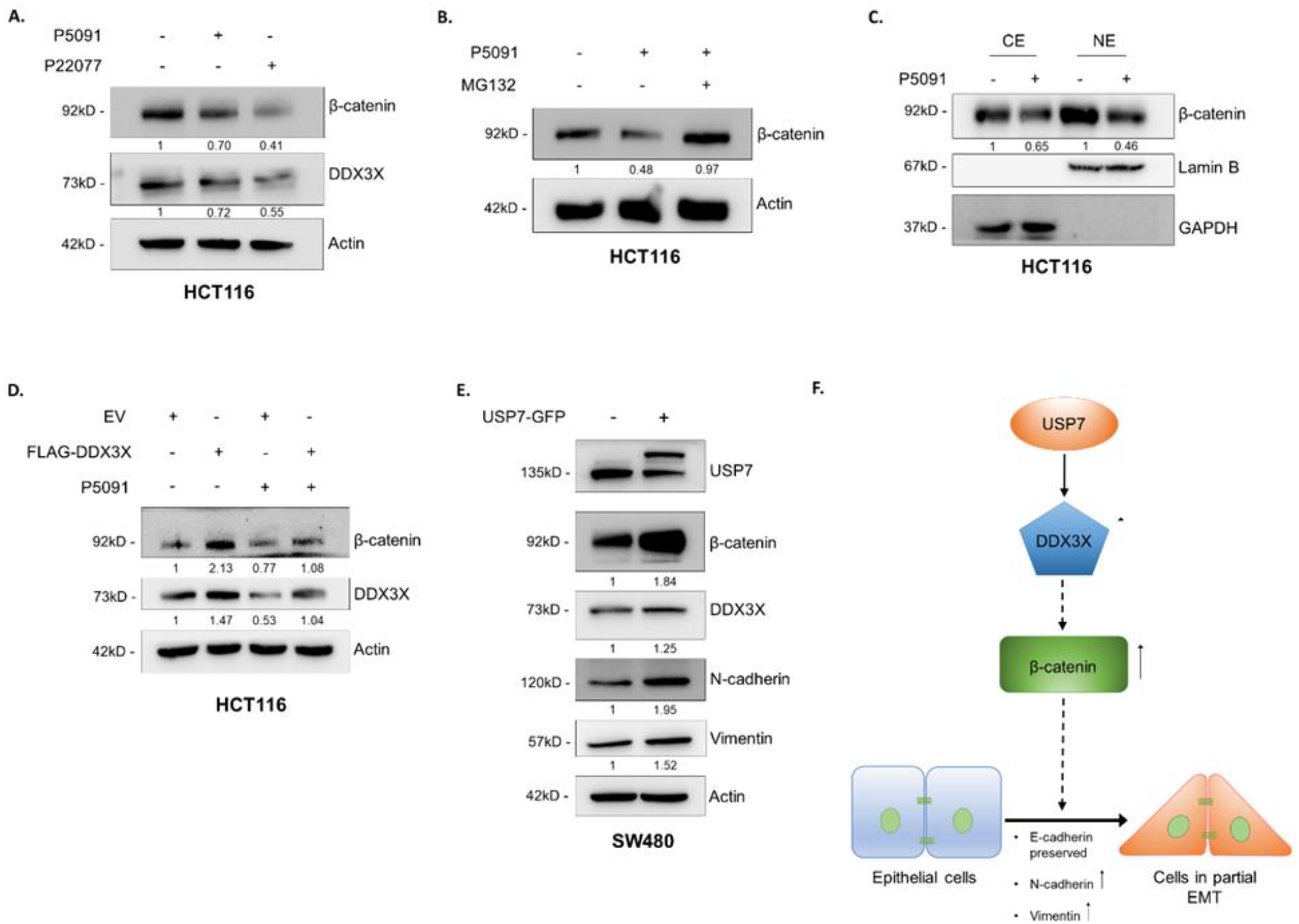
USP7 augments Wnt/β-catenin signaling through DDX3X stabilization to promote. mesenchymal phenotype. (A) HCT116 cells were treated with either DMSO control, P5091 (10 μM) or P22077 (15 μM) for 48 h. Whole cell lysates were analyzed by western blotting using the indicated primary antibodies. (B) HCT116 cells were treated with either DMSO control or P5091 (10 μM) for 24 h. 6 h prior to harvesting cells were treated with either DMSO control or MG132 (30 μM). Whole cell lysates were analyzed by western blotting using the indicated primary antibodies. (C) HCT116 cells were treated with either DMSO control or P5091 (10 μM) for 24 h. Cytosolic and nuclear fractions were analyzed by western blotting using the indicated primary antibodies. (D) HCT116 cells were transfected with either empty vector or FLAG-DDX3X as indicated. 24 h post transfection cells were either treated with DMSO control or P5091 (10 μM) as indicated for 24 h. Whole cell lysates were analyzed by western blotting using the indicated primary antibodies. (D) SW480 cells were transfected with either empty vector or GFP-USP7. Whole cell lysates were analyzed by western blotting using the indicated primary antibodies. (E) Model showing that USP7 upregulates DDX3X stability which in turn increases β-catenin levels and may lead to the development of partial EMT state. Dashed arrows signify different mechanisms of action.

Finally, we were curious to see the effects of USP7 supplementation on the mesenchymal features of colon cancer cells. As we mentioned earlier SW480 cells displayed lower endogenous levels of USP7 and showed less prominent mesenchymal features compared to HCT116 cells. Exogenous expression of USP7 in the SW480 cell line produced a marked increase in β-catenin protein levels and also in the levels of the mesenchymal markers N-cadherin and Vimentin (Fig. 6E). Taken together our data strongly indicate a positive role of USP7 in promoting Wnt/β-catenin signaling through the stabilization of DDX3X and we theorize this to be a major pathway through which USP7 controls the regulation of mesenchymal markers.

## Discussion

Till date cancer remains one of the most complex and fatal diseases to plague humanity. This complexity can be accredited to the sheer number of genetic and protein aberrations, dysfunction in major regulatory pathways, cell-environment interactions, organ of origin and even the specific sequence in which the aforementioned phenomena take place. Despite the vast amount of time and resources being pooled into cancer research our understanding of the disease remains to be incomplete.

In this study, we found that the deubiquitinase USP7 is overexpressed in CRC and its expression is associated with increasing tumor stage, thereby indicating a link between USP7 and cancer progression. Advanced stages of cancer are characterized by the induction of EMT and migration of tumor cells to adjacent tissues and lymph nodes. IHC analysis of CRC tissues revealed a strong correlation between USP7 and the mesenchymal markers N-cadherin and Vimentin. Interestingly, no significant correlation could be drawn between USP7 and the epithelial marker E-cadherin. In addition, majority of the CRC samples showed positive expression of E-cadherin. As per the classical notion of EMT, tumor cells discard their epithelial characteristics and in exchange acquire mesenchymal characters, thereby resulting in greater motility [1, 2]. According to this view, the upregulation of mesenchymal markers such as N-cadherin and Vimentin should be associated with the downregulation of epithelial markers like E-cadherin. However, our IHC analysis revealed that E-cadherin was maintained even in tumors displaying high levels of N-cadherin and Vimentin. It was possible that these results could be attributed to the small sample population of our analysis and so we decided to investigate this scenario at the *in vitro* level.

Functional inhibition of USP7 resulted in a marked reduction in migration ability of colon cancer cell lines which we assessed through 2 different migration assays. At the protein level, we observed that functional inhibition of USP7 resulted in a significant reduction in N-cadherin and Vimentin levels. In addition, E-cadherin levels did show a certain degree of upregulation in response to USP7 inactivation. However, upon silencing of USP7 expression by means of USP7-specific siRNA, we found that only the mesenchymal markers showed downregulation comparable to the results obtained during the USP7 inhibition experiments. E-cadherin levels did not show any marked changes upon silencing of USP7 expression. This was in line with the observations we made during the IHC analysis of CRC tissues. We believe that the upregulation of E-cadherin observed in response to inactivation of USP7 was most likely due to off-targets effects of the USP7 inhibitors P5091 and P22077.

Contrary to the classical view that loss of E-cadherin expression is essential for EMT induced cancer cell migration, recent studies have indicated that this may not always be the case. It has been demonstrated that EMT is not an all-or-none process i.e., a complete switch from epithelial to mesenchymal phenotype, and instead represents a multi-stage process consisting of hybrid epithelial-mesenchymal states in between completely epithelial and completely mesenchymal phenotypes. These hybrid states are generally termed EMT-like or partial EMT and exhibit the expression of both epithelial and mesenchymal markers [7, 8]. Many aggressive and metastatic tumors have been shown to retain E-cadherin expression on their surface [41–43]. One particular study carried out on a wide variety of cancer types including CRC has shown that compared to single cell dissemination and migration from the invasive front of the tumor, the dissemination and migration of cell clusters is far more common, which in turn indicates the persistence of certain amount of epithelial cell-cell adhesions [44]. Apart from this, there are studies that have demonstrated that the presence of E-cadherin may have a positive effect on the dissemination, migration and survival of cancer cells [45, 46]. Our data indicates that USP7 positively regulates N-cadherin and Vimentin levels, thereby promoting the acquisition of mesenchymal characters. However, USP7 showed no significant effect on E-cadherin levels. From this we theorized that USP7 promotes the development of a partial EMT state in colon cancer cells. It is to be noted that only the total intracellular levels of E-cadherin protein were assessed in this study and not just plasma membrane associated E-cadherin.

Through the utilization of a proteomics-based approach we identified the RNA helicase DDX3X to be a possible interacting partner of USP7 which we then went on to verify *in vitro*. We further show that binding of USP7 to DDX3X is facilitated by the TRAF domain of USP7. Experiments involving the inhibition of USP7 catalytic activity as well as the overexpression of both wild type USP7 and its catalytic mutant revealed that USP7 stabilizes DDX3X *via* its deubiquitinase activity. This strongly suggests that USP7 can regulate the functions associated with DDX3X. As DDX3X has previously been shown to be involved in promoting tumor metastasis in cancer cells including colon cancer cells, there was a strong probability that the observed downregulation of mesenchymal characters in colon cancer cells upon USP7 inhibition or depletion was due to the lack of stabilization and subsequent loss of DDX3X. In accordance with previous studies, we found that downregulation of DDX3X upon inhibition of USP7 deubiquitinase activity resulted in a comparable reduction in β-catenin levels. We also demonstrated that β-catenin levels could be rescued upon exogenous supplementation of DDX3X. Lastly, we showed that supplementation of USP7 in colon cancer cell lines that displayed lower endogenous levels of USP7 resulted in the upregulation of β-catenin as well as the mesenchymal markers N-cadherin and Vimentin. Wnt/β-catenin signaling is generally considered a major oncogenic signaling pathway involved in the induction of EMT through the regulation of different core EMT transcription factors [1, 5]. The transcription factor OVOL has been shown to be essential for the stabilization of partial EMT phenotype and silencing of OVOL gene expression is associated with transition to complete EMT [47, 48]. Interestingly, OVOL is a downstream target of Wnt/β-catenin signaling [49, 50]. It is possible that enhanced Wnt/β-catenin signaling induced by the stabilization of DDX3X by USP7 may lead to upregulation of OVOL thereby preventing the complete transition to EMT and instead allowing for a partial EMT. It is possible that additional oncogenic events are required for the final transition to complete EMT. Further investigations into the mechanistic details underlying partial EMT induction, stability and transitioning to complete EMT are required.

In conclusion, the findings of this investigation indicate a positive role of USP7 in controlling the acquisition and maintenance of mesenchymal phenotype in CRC. We report here that the RNA helicase DDX3X is a novel interacting partner of USP7 and that USP7 mediates the deubiquitination and stabilization of DDX3X. We also report that USP7 mediated stabilization of DDX3X leads to upregulation of β-catenin levels in colon cancer cell lines. This finding is supported by previous studies which have reported a pro-regulatory role of USP7 in Wnt/ β-catenin signaling [37–39]. We further report that USP7 upregulation does not elicit a complete EMT response and instead promotes the development of a partial EMT state. Taken together, this study highlights the importance of USP7 in CRC progression and the need to place it as a high priority target during the designing of future therapeutic approaches aimed at curbing CRC.

## Acknowledgments

Authors sincerely acknowledge Dr. Seemana Bhattacharya and Dr. Gouranga Saha (ex-students of Dr. Mrinal K Ghosh) for their technical help in developing this project and current findings.

## Data Availability

Data supporting the conclusion of the current study is included in the manuscript and supplementary files.

