## Supplementary file for "A novel role of USP7 in imparting partial EMT state in colorectal cancer through the DDX3X-β-catenin axis"

Figure S1

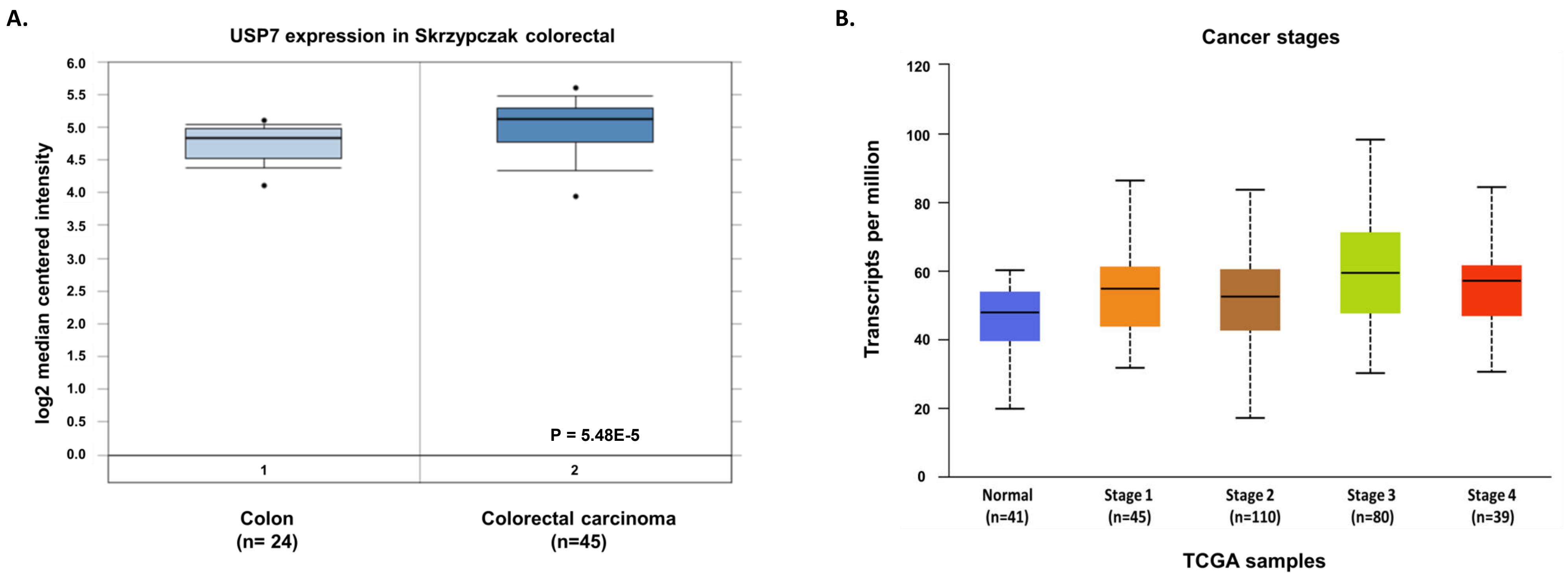

A.
USP7 expression in Skrzypczak colorectal
log2 median centered intensity
6.0
5.5
5.0
4.5
4.0
3.5
3.0
2.5
2.0
1.5
1.0
0.5
0.0
P = 5.48E-5
1
2
Colon
(n= 24)
Colorectal carcinoma
(n=45)
B.
Cancer stages
Transcripts per million
120
100
80
60
40
20
0
Normal
(n=41)
Stage 1
(n=45)
Stage 2
(n=110)
Stage 3
(n=80)
Stage 4
(n=39)
TCGA samples

**Figure S2**

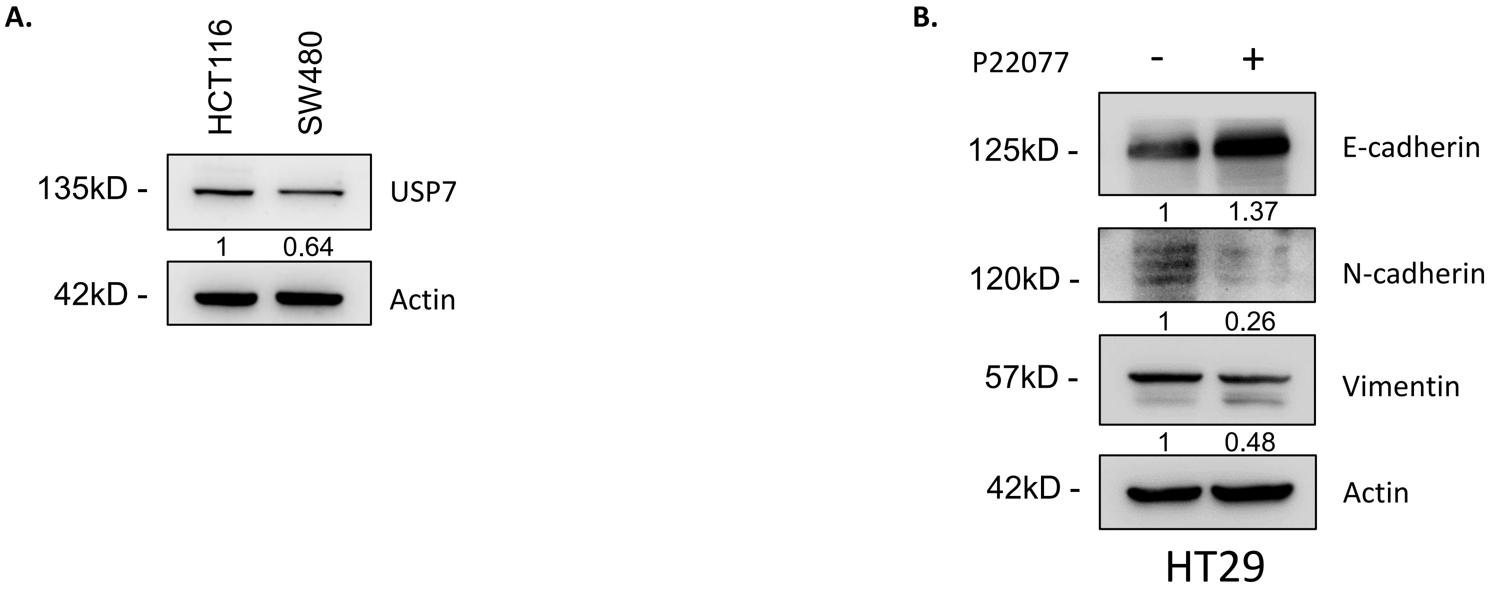

A.
HCT116
SW480
135kD -
USP7
1
0.64
42kD -
Actin
B.
P22077
-
+
125kD -
E-cadherin
1
1.37
120kD -
N-cadherin
1
0.26
57kD -
Vimentin
1
0.48
42kD -
Actin
HT29

**Figure S3**

**A.**

**DDX3X**

MSHVAVENALGLDQQFAGLDLNSSDNQSGGSTASKGRYIPPHLRNREATKGFYDKDSSGWSSSKDK
DAYSSFGSRSDSRGKSSFFSDRGSGSRGRFDDRGRSDYDGIGSRGDRSGFGKFERGGNSRWCDKSD
EDDWSKPLPPSERLEQELFSGGNTGINFEKYDDIPVEATGNNCPPHIESFSDVEMGEIIMGNIELT
RYTRPTPVQKHAIPIIKEKRDLMACAQTGSGKTAAFLL**PILS**QIYSDGPGEALRAMKENGRYGRR
KQYPISLVLAPTRELAVQIYEEARKFSYRSRVRPCVVYGGADIGQQIRDLERGCHLLVATPGRLVD
MMERGKIGLDFCKYLVLDEADRMLDMGFEPQIRRIVEQDTMPPKGVRHTMMFSATFPKEIQMLARD
FLDEYIFLAVGRVGSTSENITQKVVWVEESDKRSFLLDLLNATGKDSLTLVFVETKKGADSLEDFL
YHEGY**ACTS**IHGDRSQRDREEALHQFRSGKSPILVATAVAARGLDISNVKHVINFDLPSDIEEYV
HRIGRTGRVGNLGLATSFFNERNINITKDLLDLLVEAKQEVPSWLENMAYEHHYKGSSRGRSKSSR
FSGGFGARDYRQSSG**ASSS**SFSSSR**ASSS**RSGGGGHGSSRGFGGGGYGGFYNSDGYGGNYNSQGVD
WWGN

**B.**

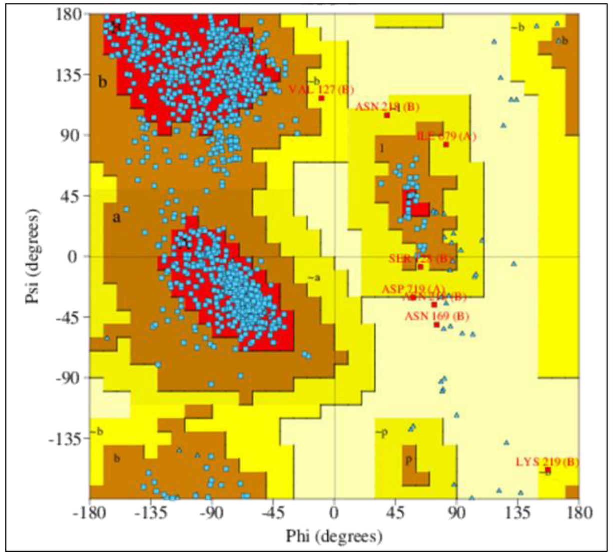


**C.**

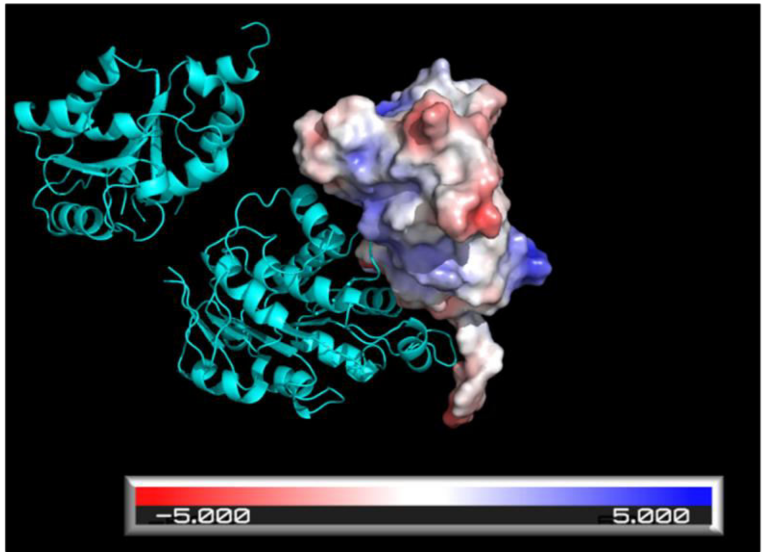


**D.**

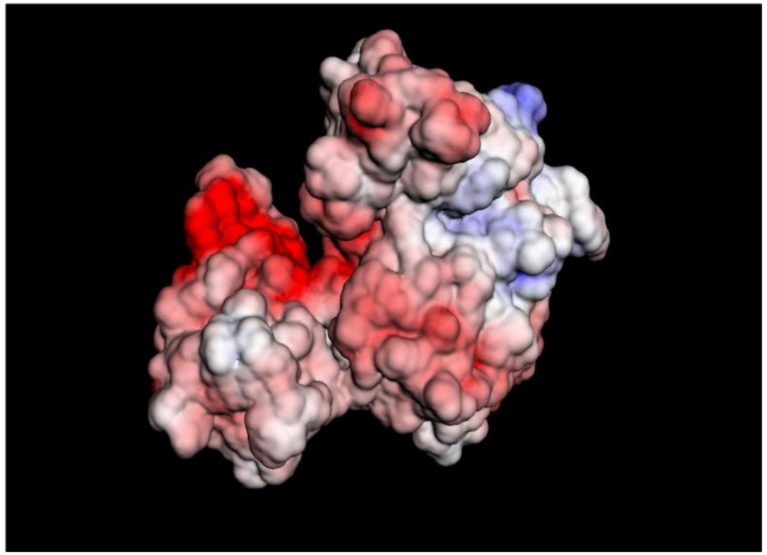

### SUPPLEMENTARY FIGURE LEGENDS

**Figure S1.** (A) USP7 gene expression in normal colon vs colon carcinoma tissues from the Skrzypczak colorectal dataset. (B) Cancer stage specific (I-IV) study of USP7 expression level of tumor samples in comparison with normal tissue samples in UALCAN analysis.

**Figure S2.** (A) HT29 cells were treated with either DMSO control or P5091 (10 μM) for 48 h. Whole cell lysates were analyzed by western blotting using the indicated primary antibodies. (B) Whole cell lysates of HCT116 and SW480 CELLS were analyzed by western blotting using s primary antibodies against USP7.

**Figure S3.** (A) Complete amino acid sequence of DDX3X. Sequences in bold font represent possible target sequences for binding with USP7 TRAF domain. (B) Evaluation of stereochemical quality of the docked structure showed that 81.9% of residues lay in the favored region of Ramachandran plot. (C) Adaptive Poisson–Boltzmann Solver (APBS) electrostatic interactions of docked structure. (D) Predicted surface potential of docked structure. Negatively and positively charged surface areas are depicted in red and blue, respectively.
